# Imaging voltage in complete neuronal networks within patterned microislands reveals preferential wiring of excitatory hippocampal neurons

**DOI:** 10.1101/2020.10.09.332304

**Authors:** Alison S. Walker, Benjamin K. Raliski, Dat Vinh Nguyen, Patrick Zhang, Kate Sanders, Kaveh Karbasi, Evan W. Miller

## Abstract

Voltage imaging with fluorescent dyes affords the opportunity to map neuronal activity in both time and space. One limitation to imaging is the inability to image complete neuronal networks: some fraction of cells remains outside of the observation window. Here, we combine voltage imaging, *post hoc* immunocytochemistry, and patterned microisland hippocampal culture to provide imaging of complete neuronal networks. The patterned microislands completely fill the field of view of our high-speed (500 Hz) camera, enabling reconstruction of the spiking patterns of every single neuron in the network. Cultures raised on microislands develop similarly to neurons grown on coverslips and display similar composition of inhibitory and excitatory cell types. The principal excitatory cell types (CA1, CA3, and dentate granule cells, or DGC) are also present in similar proportions in both preparations. We calculate the likelihood that action potential firing in one neuron to trigger action potential firing in a downstream neuron in a spontaneously active network to construct a functional connection map of these neuronal ensembles. Importantly, this functional map indicates preferential connectivity between DGC and CA3 neurons and between CA3 and CA1 neurons, mimicking the neuronal circuitry of the intact hippocampus. We envision that patterned microislands, in combination with voltage imaging and methods to classify cell types, will be a powerful method for exploring neuronal function in both healthy and disease states. Additionally, because the entire neuronal network is sampled simultaneously, this strategy has the power to go further, revealing all functional connections between all cell types.

**Significance Statement:** *In vitro* model systems provide unsurpassed control and access for exploring the molecular and cellular details of neurobiology. We developed a patterned microisland system for culturing rat hippocampal neurons that recapitulates the features of bulk hippocampal cultures, but with the added benefit of allowing access to high-speed imaging of entire neuronal ensembles using voltage imaging. By using far-red voltage-sensitive fluorophores, we map the functional connections across all cells in the neuronal ensemble, revealing that several important functional synapses present in the intact hippocampus are recapitulated in this microisland system. We envision the methods described here will be a powerful complement to ongoing research into basic neurobiological mechanisms and the search for therapies to treat diseases arising from their dysfunction.

## Introduction

Neurons, the electrically excitable cells of the nervous system, are organized together in circuits that perform specific functions: from detecting motion across our retinas to storing memories and higher cognition. The function of circuits is determined by intrinsic morphological, chemical, and biophysical properties of the neuronal cell types included, as well as the number, strength, and placement of connections between them. The complexity of all these components means that understanding neuronal circuits remains one of the most challenging questions in modern day neuroscience.

To start to understand how the brain functions at the level of synapses and circuits, we need to know what cell types are present and how they wire up to each other in microcircuits. The field of “Connectomics” attempts to unravel this complexity using transmission electron microscopy to painstakingly map pathways between neurons and their connections. However, these analyses are laborious, with a recent reconstruction of a single cubic millimeter of mouse brain, taking 8 months to complete and generating 2 million gigabytes of data. Additionally, these connectomes are structural in nature. The presence of synaptic connections cannot tell us how often they are used and whether their activation would lead to forward transmission of information, i.e. an action potential in the receiving neuron.

Recent technological advances in calcium imaging have led to fascinating observations of neural correlates of sensation and behavior (Chen et al., 2013; Vladimirov et al., 2014; Barson et al., 2020). However, precise neuronal wiring is blurred by the slow decay of calcium signals, meaning large proportions of cells or brain regions appear simultaneously ‘on’ without the resolution to tease apart temporal triggering of connected single cells. Another strategy to probe the function of circuits is to activate neurons with either electrical stimulation or optogenetic activation, recording the effect on single postsynaptic neurons electrophysiologically. While highly detailed, extrapolating findings from very small numbers of recorded neurons leads to generalizations of complex interactions of multiple neurons.

To complement the understanding of neuronal wiring specificity and neuronal circuitry, here we have developed a voltage imaging approach to reveal neuronal wiring in dissociated hippocampal microcircuits. The tractability of reductionist *in vitro* models makes them an excellent starting point to probe cellular and molecular players that are essential to the development and health of neural circuits *in vivo* (Turrigiano et al., 1998; Burrone et al., 2002; Williams et al., 2011). In addition, *in vitro* assays and methods are central to phenotypic screening efforts to uncover new drug candidates and druggable targets. To ensure *in vitro* screening data translates well *in vivo*, and ultimately to clinical settings, increased emphasis is placed on a deeper understanding of these models. Hippocampal culture models are often used to study neuronal activity due to the importance of the hippocampus in learning and memory (Bird and Burgess, 2008), as well as a site of seizure generation (Chatzikonstantinou, 2014). However, it is not known how completely the functional connectivity of hippocampal neurons is conserved in monolayer culture *in vitro*.

Here we outline and validate patterning techniques to generate functional, isolated microislands of up to 25 neurons. Patterned microislands of defined size enable voltage imaging of a complete neuronal network, with all possible connections observable within the imaging window. We show that these reductionist models conserve the same functional properties as cultures 1000x larger and can be used to match functional signatures to cell type identity. Building on this, we use the timing of spontaneous action potential firing to infer functional connectivity, i.e. how likely a neuron is to trigger action potential firing in a connected neuron. We pair this probability map with a cell type atlas to demonstrate that preferential wiring of hippocampal neurons *in vivo* is conserved in microisland culture *in vitro*. Together, this work introduces a means to profile neuron-to-neuron wiring in an accessible system amenable to study microcircuit behavior in development, health, and disease.

## Materials and Methods

### Cell Culture

All animal procedures were approved by the UC Berkeley Animal Care and Use Committees and conformed to the NIH Guide for the Care and Use of Laboratory Animals and the Public Health Policy.

### Microislands

Generation of PDMS stencils was performed as previously reported (Hoang et al., 2018). An SU-8 master was fabricated by spin coating a 4-inch silicon water (University Wafer, TX) with SU-8 50 negative photoresist. A photomask (CAD/Art Services, http://www.outputcity.com/) with opaque regions equal to the size of our imaging window (650 μm x 120 μm) was placed on top to the photoresist coated wafer and exposed to UV light. Following development, the final silicon wafer was fabricated with pillars equal to our imaging size. This work was performed in the Biomolecular Nanotechnology Center (BNC) at the University of California, Berkeley. This silicon wafer was used to prepare 4-polydimethylsiloxane (PDMS) stencils and was silianzed with trimethylchlorosilane once every 4 PDMS coatings to prevent PDMS bonding to the wafer. PDMS stencils were prepared by pouring a PDMS mixture (SYLGARD 184; 10:1 monomer:catalyst) onto the silicon wafer and placing the wafer between two plastic transparencies followed by two pieces of glass. This assembly was held together with clips and cured in a 65 °C oven for 2 hours. The assembly was then disassembled and the PDMS stencil was removed from the silicon wafer. The stencil sheet was then cut into smaller stencils containing 4 × 8 holes and these smaller stencils were stored in 70% ethanol indefinitely. These stencils were then placed on 12 mm diameter coverslips (Fisher Scientific) on the lid of a 24-well cell culture plate (VWR) and allowed to dry for at least 2 hours. The coverslips were then etched with oxygen plasma with a reactive-ion etching (RIE) system in the Biomolecular Nanotechnology Center (BNC) at the University of California, Berkeley and stored in a 24-well cell culture plate in ethanol at 4 °C until needed. These coverslips were washed with 70% ethanol, followed by DPBS (Gibco) and used to prepare cultures of dissociated rat hippocampal neurons (see below). At 4-7 DIV, the stencils were removed from the coverslips using ethanol sterilized forceps. Stencils were removed at least 2 days prior to functional imaging.

### Rat Hippocampal Neurons

Hippocampi were dissected from embryonic day 18 Sprague Dawley rats (Charles River Laboratory) in cold sterile HBSS (zero Ca^2+^, zero Mg^2+^). All dissection products were supplied by Invitrogen, unless otherwise stated. Hippocampal tissue was treated with trypsin (2.5%) for 15 min at 37 °C. The tissue was triturated using fire polished Pasteur pipettes, in minimum essential media (MEM) supplemented with 5% fetal bovine serum (FBS; Thermo Scientific), 2% B-27, 2% 1 M D-glucose (Fisher Scientific) and 1% GlutaMax. The dissociated cells were plated onto 12 mm diameter coverslips (Fisher Scientific) for bulk culture or with an affixed PDMS stencil (as outlined above) for microisland culture and pre-treated with PDL. Cells were plated at a density of 30-40,000 cells per coverslip in MEM supplemented media. For experiments detailed neuronal subtype specific firing rates in large cultures, cells were plated onto 3 cm gridded dishes (ibidi) at a density of 125k/dish. Neurons were maintained at 37 °C in a humidified incubator with 5% CO_2_. At 1 day in vitro (DIV), half of the MEM supplemented media was removed and replaced with Neurobasal media containing 2% B-27 supplement and 1% GlutaMax. Functional imaging was performed on 8-15 DIV neurons to access neuronal excitability and connectivity across different stages of development.

### BeRST 1 Stocks and Cellular Loading

For all imaging experiments, BeRST 1 was diluted from a 250 μM DMSO stock solution to 0.5 μM in HBSS (+Ca^2+^, +Mg^2+^, -phenol red). To load cells with dye solution, the media was first removed from a coverslip and then replaced with the BeRST-HBSS solution. The dye was then allowed to load onto the cells for 20 minutes at 37 °C in a humidified incubator with 5% CO_2_. After dye loading, coverslips were removed from the incubator and placed into an Attofluor cell chamber filled with fresh HBSS for functional imaging. The location of imaged islands within the 4 × 8 microisland grid was noted to aid reloaction following post hoc immunocytochemistry.

### Immunocytochemistry – Cell Type Identity and Synaptic Development

After imaging, neurons were fixed by placing the coverslips in a 4% paraformaldehyde (PFA) solution for 20 minutes and then washing 3x with PBS (for cell type identity analysis). For synaptic development analysis, coverslips of neurons were fixed by treatment for 10 minutes in a 50:50 mixture of ice-cold methanol (MeOH) and PBS, followed by 10 minutes in 100% ice-cold MeOH. Coverslips were washed 3x with PBS. Coverslips were then stored in PBS at 4 °C for up to a week before carrying out the following immunostaining protocol. Briefly, neurons were permeabilized with 0.25% Triton-X in PBS for 5 minutes. The neurons were then washed 3x with PBS and blocked with 10% Native Goat Serum (NGS) in PBS for 2 hours on a shaker at room temperature. Primary antibodies were diluted to the concentrations indicated below in PBS with 2% NGS. The 10% NGS in PBS solution was aspirated off of the neurons and replaced with the primary antibody solutions. The neurons were then placed on a shaker at room temperature for 2 hours. After 2 hours, the neurons were washed 5x with PBS, loaded with secondary antibodies diluted to the concentrations indicated below in PBS with 2% NGS, and placed on a shaker in the dark for 45 minutes at room temperature. Finally, the neurons were washed 5x with PBS and re-imaged following placement in an Attofluor cell chamber.

### Primary Antibodies – Cell Type identity

Antibody (Supplier, Dilution) – Neuronal Cell Type Labeled

Mouse IgG1 anti-CaMKII (Millipore EMD, 1:2000) – Excitatory neurons

Mouse IgG2b anti-Prox1 (Sigma-Aldrich, 1:2000) – Dentate Granule cells

Mouse IgG2a anti-GAD67 (Millipore EMD, 1:1500) – Inhibitory (GABAergic) neurons

Rat IgG anti-CTIP2 (Abcam, 1:500) – CA1 neurons

### Secondary Antibodies

Antibody (Supplier, Dilution) – Fluorescent Dye

Goat anti-mouse IgG1 (Biotium, 1:2000) – CF405S

Goat anti-mouse IgG2b (Invitrogen, 1:2000) – Alexa Fluor 488

Goat anti-mouse IgG2a (Life Technologies, 1:2000) – Alexa Fluor 546

Goat anti-rat IgG (Life Technologies, 1:2000) – Alexa Fluor 647

### Primary Antibodies for Synaptic Development

Rabbit IgG anti-Synapsin 1 – (Synaptic Systems, 106103)

Mouse IgG2aκ anti-PSD-95 – (Millipore EMD MABN68, clone K28/43)

Rabbit IgG anti-MAP2 – (Abcam, ab32454)

Hoescht 33342

### Imaging Parameters

#### Spontaneous Activity Imaging

Spontaneous activity imaging was performed on an upright AxioExaminer Z-1 (Zeiss) or an inverted Zeiss AxioObserver Z-1 (Zeiss), both equipped with a Spectra-X light engine LED light (Lumencor), and controlled with μManager (V1.4, open-source, Open Imaging) (Edelstein et al., 2014). Images were acquired using a W-Plan-Apo/1.0 NA 20x water immersion objective (Zeiss) or a Plan-Apochromat/0.8 NA 20x air objective (Zeiss). Images (2048 x 400 px^2^, pixel size: 0.325 x 0.325 μm^2^) were collected continuously on an OrcaFlash4.0 sCMOS camera (sCMOS; Hamamatsu) at a sampling rate of 0.5 kHz, with 4×4 binning, and a 633 nm LED excitation light power of 6.5 mW/mm^2^. 6 x 10 s movies were recorded sequentially for each imaging area.

#### Immunofluorescence

Cell-type immunofluorescence was recorded on an upright AxioExaminer Z-1 (Zeiss) or an inverted Zeiss AxioObserver Z-1 (Zeiss), both equipped with a Spectra-X light engine LED light (Lumencor), and controlled with Slidebook software (Intelligent Imaging Innovations). Neuronal cell type was determined by the presence or absence of cell type specific proteins as detected by immunofluorescence. All neurons with GAD67 present were labeled as inhibitory (GABAergic) neurons. All neurons with CAMKII plus Prox1 staining were labeled as dentate granule cells. All neurons with CaMKII and CTIP2 present were labelled as CA1 neurons. All neurons with CaMKII present but neither CTIP2 nor Prox 1 were labelled as CA3 neurons.

Synaptic development immunofluorescence was recorded using an upright Zeiss LSM 880 confocal (Zeiss) and controlled with Zen2 software (Zeiss). Samples were imaged with a 40x objective.

#### Image Analysis

All imaging analysis was performed using SpikeConnect, a MATLAB script developed in-house to detect action potentials from fluorescence traces associated with manually drawn regions of interest. A brief description of the SpikeConnect workflow is outlined below. This code is available from GitHub upon request. The code was added to the MATLAB path and can be run un on all versions post 2017a. The imaging data was organized in the following manner. Each coverslip was organized into individual folders which contained separate folders for each area imaged on that coverslip. Each area folder contained a brightfield image of the area and the fluorescence movies recorded from that area. All plotting and statistical analysis was performed in GraphPad.

#### Drawing Regions of Interest (ROIs) and Labeling Neurons

Individual neurons were labeled by running “selectroi_gui”, selecting a brightfield image (as a .tiff file) and importing the movies associated with the image (also as .tiff files). The frame rate parameter was set to the recording frame rate. Regions of interest (ROIs) were drawn around each neuron in the brightfield image and saved. Next, a ROI was drawn around an area of background in the first frame of the fluorescence movie associated with the brightfield image and saved. This process was repeated for each area of imaging. If cell type information was known from correlative immunostaining experiments, neuron ROIs were labeled as DGC neurons, inhibitory neurons, CA1 neurons, CA3 neurons, or unknown by running “labelroi”.

#### Action Potential Detection

Action potentials (spikes) were detected by running “batchkmeans_gui”, selecting area folders for analysis, and using the background ROI to generate background-corrected fluorescence traces for each neuron ROI. This script then uses k-means clustering to identify possible action potentials (spikes), subthreshold events, and baseline. After the running the k-means clustering algorithm, a signal-to-noise ratio (SNR) threshold for a signal to be labeled as a spike was set by running “thresholding_gui”. This threshold was established from dual electrophysiology-optical experiments as the optical trace threshold which faithfully reproduces the spikes detected by electrophysiology. The spike data was then saved for further analysis.

#### Firing Frequency Analysis

Firing frequencies from the spike data were exported as excel files for each ROI by running “freqexport_gui”. The excel files generated contained average frequency (Hz), instantaneous frequency (Hz), and interspike interval (ms) data for each ROI along with summary columns for each of these three parameters. This data was then plotted in GraphPad for comparison.

#### Spike Time Tiling Coefficient (STTC) Analysis

Spike time tiling coefficients (STTC) are the correlation between a pair of spike trains. These were calculated by selecting spike data and running “sttc_gui”. The resulting STTC values for each ROI pair were then exported as an excel file and plotted in GraphPad for comparison.

##### XCI Analysis

The metric XCI between neurons 1 and 2 is the fraction of 1’s spikes that have a corresponding spike in 2’s spike train within a certain defined lag time period. This metric was calculated by selecting spike data and running “xci_gui”. The connectivity factor summaries were exported as excel files and plotted in GraphPad for comparison.

#### Generation of Image Sequence in Figure 1i and Movie S1

**Figure 1.**
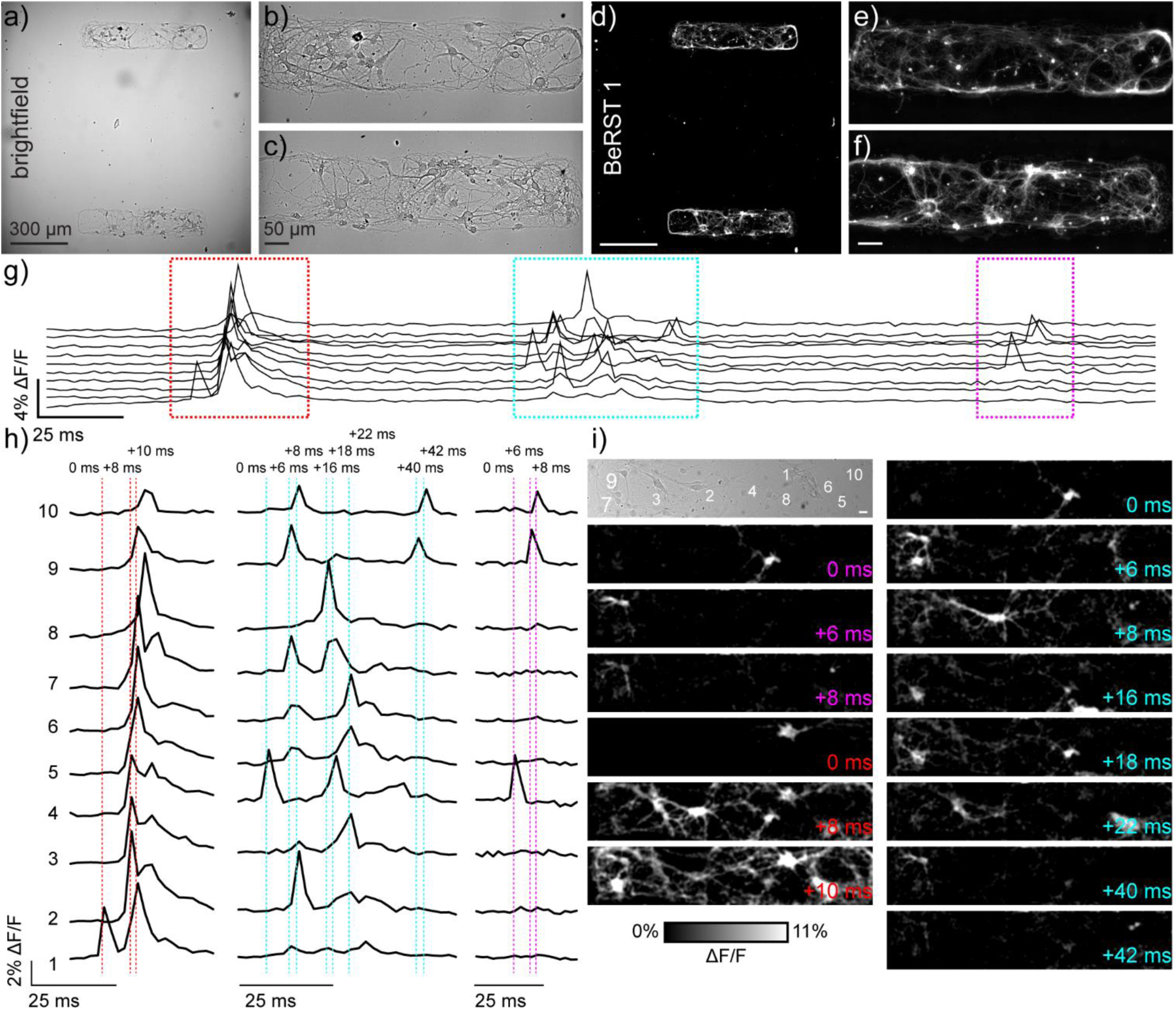
Monitoring neuronal activity in patterned microislands of dissociated rat hippocampal neurons using BeRST 1. **(a)** Representative differential interference contrast (DIC) image of dissociated rat hippocampal neurons grown on patterned microislands. Higher magnification image of **(b)** upper and **(c)** lower microisland from panel **(a)**. **(d-f)** Corresponding BeRST 1 (500 nM) fluorescence images. **(g)** Plot of ΔF/F vs time for a representative microisland culture stained with BeRST 1 and imaged at 500 Hz. **(h)** Expanded times scale for colored, boxed regions in panel (g). Dashed vertical lines and timing labels indicate relative timing of fluorescence intensity changes. **(i)** Transmitted light image of neurons recorded in panels (g) and (h), with numbers indicating the neurons plotted in (g) and (h). Images with colored time stamps are ΔF/F images corresponding to single frames (2 ms exposure) from the recorded image sequence at the dashed vertical lines in panel (h). Images have a 2 pixel Gaussian blur applied to aid visualization. Scale bar for (i) is 20 μm.

We imported the 16-bit grey value image sequence into FIJI (v1.52g). The image stack was stabilized using the StackReg>RigidBody plugin. We created a ΔF movie by first creating an Average Z-Projection from images 1-25 (Stack>ZProjection>Average) and then subtracting this image from each image in the sequence. This new ΔF movie (32-bit) was then divided by the same Average Z-Projection image to create a ΔF/F movie (32-bit). We scaled the histrogram values from 0 to 11 percent ΔF/F and converted to an 8 bit image. We applied a 2 pixel Gaussian blur to each frame (Process>Filter>Gaussian Blur) to smooth out noise and aid visibility. Individual frames were included in the image sequence in Figure 1i. The entire image stack was converted to an AVI file (no compression, 25 fps playback) and included as Movie S1.

## Results

### Utilizing microcontact printing to enable circuit-wide interrogation by voltage imaging

Voltage imaging is a powerful technique which can be used to simultaneously record the functional signatures of multiple neurons with high sensitivity and temporal resolution (Huang et al., 2015; Liu and Miller, 2020; Walker et al., 2020). Previously, we recorded small areas of neuronal networks cultured on 12 mm diameter coverslips (Walker et al., 2020). The imaging field of view (FOV) comprised an area of approximately 130 × 665 μm^2^, or 0.086 mm^2^. This means that more than 99.9% of neurons on the coverslip, with a total area of 113 mm^2^, were outside our field of view, invisible to our analysis. In this imaging configuration, a substantial fraction of the inputs to and outputs from recorded neurons remains completely unknowable. Assigning neuron to neuron connectivity without direct observation of the majority of neurons in the sample is challenging. It is theoretically possible however, to use voltage imaging to examine the activity and connectivity of every neuron within a circuit empirically. To test this hypothesis, we developed a method to culture dissociated hippocampal neurons in small microislands of prescribed dimensions, where activity signatures from every single neuron could be sampled using voltage imaging.

To resolve action potentials with similar confidence to electrophysiological patch-clamp techniques, we use the far-red voltage-sensitive fluorophore, BeRST 1 (<u>Be</u>rkeley <u>R</u>ed <u>S</u>ensor of <u>T</u>ransmembrane potential) to record changes in membrane potential at an optical sampling rate of 500 Hz (Huang et al., 2015; Walker et al., 2020). For voltage imaging with sCMOS cameras, a trade-off exists between frame rate and imaging area: faster frame rates can be achieved at the expense of imaging area, while larger imaging areas are accessible only at slower frame rate. We selected 500 Hz as our optical sampling rate; it provides a balance between speeds sufficient to capture fast action potentials and provides a field of view (FOV) of approximately 120 × 650 μm^2^ (**Figure 1a-c**). To limit neuronal growth to this exact window size, we modified a lithographic technique (Hoang et al., 2018) to create polydimethylsiloxane (PDMS) stencils where the neuronal substrate, poly-D-lysine (PDL), can only access the underlying glass coverslip via holes in the stencil created to match dimensions of interest (**Figure 1-1**). This is the first time stencil patterning has been applied to neuronal culture. While the original protocol included a polyethyleneglycol (PEG) polymer film to discourage cell growth onto uncoated glass areas following stencil removal, use of a PEG film was unnecessary here because neurons do not grow well on uncoated glass. Neurons plated onto micro-patterned coverslips only developed within PDL coated areas, with numbers of neurons per microisland ranging from 10 to 25 per microisland (**Figure 1a-f**).

**Figure 1-1.**
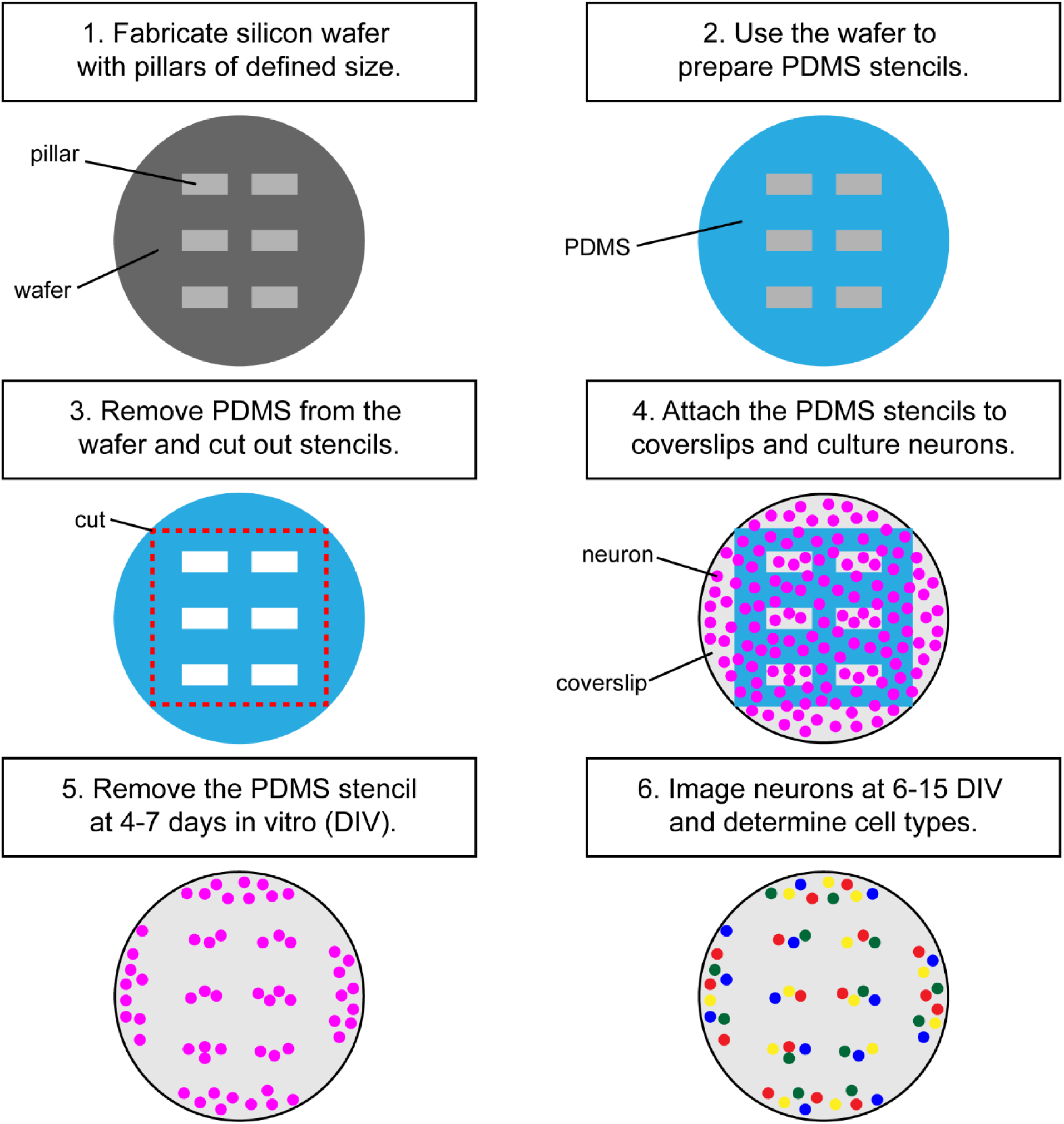
Microisland patterning workflow. These diagrams are simplified for clarity. Typically, one Polydimethylsiloxane (PDMS) sheet yields 10 stencils containing 4 × 8 microislands. Typically, microislands contain 10-20 neurons.

We used our previously established voltage imaging methodology with BeRST 1 (Walker et al., 2020) to make multiple, sequential recordings of neuronal activity from these patterned microislands (**Figure 1g**). We previously showed that low concentrations of BeRST (500 nM) and low-power, red LED illumination (<13 mW/mm^2^) enabled sustained, repeated measurements of spontaneous neuronal activity. In a typical experiment, 10-25 neurons could be observed simultaneously at an optical sampling rate of 500 Hz (**Figure 1g and h**). The fast frame rate (500 Hz) allows spike detection with 2 ms resolution (**Figure 1h and i)** and allows us to resolve bouts of activity patterns across neurons that last only 10 to 50 ms (**Figure 1 h and i**). Traditional methods of recording neuronal activity, like whole-cell patch-clamp electrophysiology requires elaborate instrumentation to record from 10 cells simultaneously (Perin and Markram, 2013; Wang et al., 2015). Imaging approaches that rely on Ca^2+^ signals would be similarly unable to parse these fast signals: the off-rate, or decay kinetics, of Ca^2+^ unbinding from the widely used genetically encoded Ca^2+^ indicator, GCaMP6f, is approximately 200 – 400 ms (Chen et al., 2013), precluding its use for resolving rapid spike trains that persist for <50 ms (**Figure 1g-i**, **Movie S1**) across multiple cells.

### Neurons cultured on microislands are functionally comparable to neurons cultured on coverslips

To determine whether neurons cultured on microislands develop similarly to *in vitro* cultures grown on coverslips (or bulk culture), we performed immunocytochemical and functional voltage imaging analyses. Qualitative immunocytochemical comparison of synaptic markers between 4 and 14 days in vitro (DIV) for neurons grown on either microislands or 12 mm coverslips (bulk culture) revealed typical patterns of synaptic markers for both network sizes (**Figure 2a**). We observe formation of presynaptic terminals (anti-Synapsin-1) prior to the appearance of excitatory postsynapses (anti-PSD-95), followed by an increase in apposition of these two markers with maturation (**Figure 2b**).

**Figure 2.**
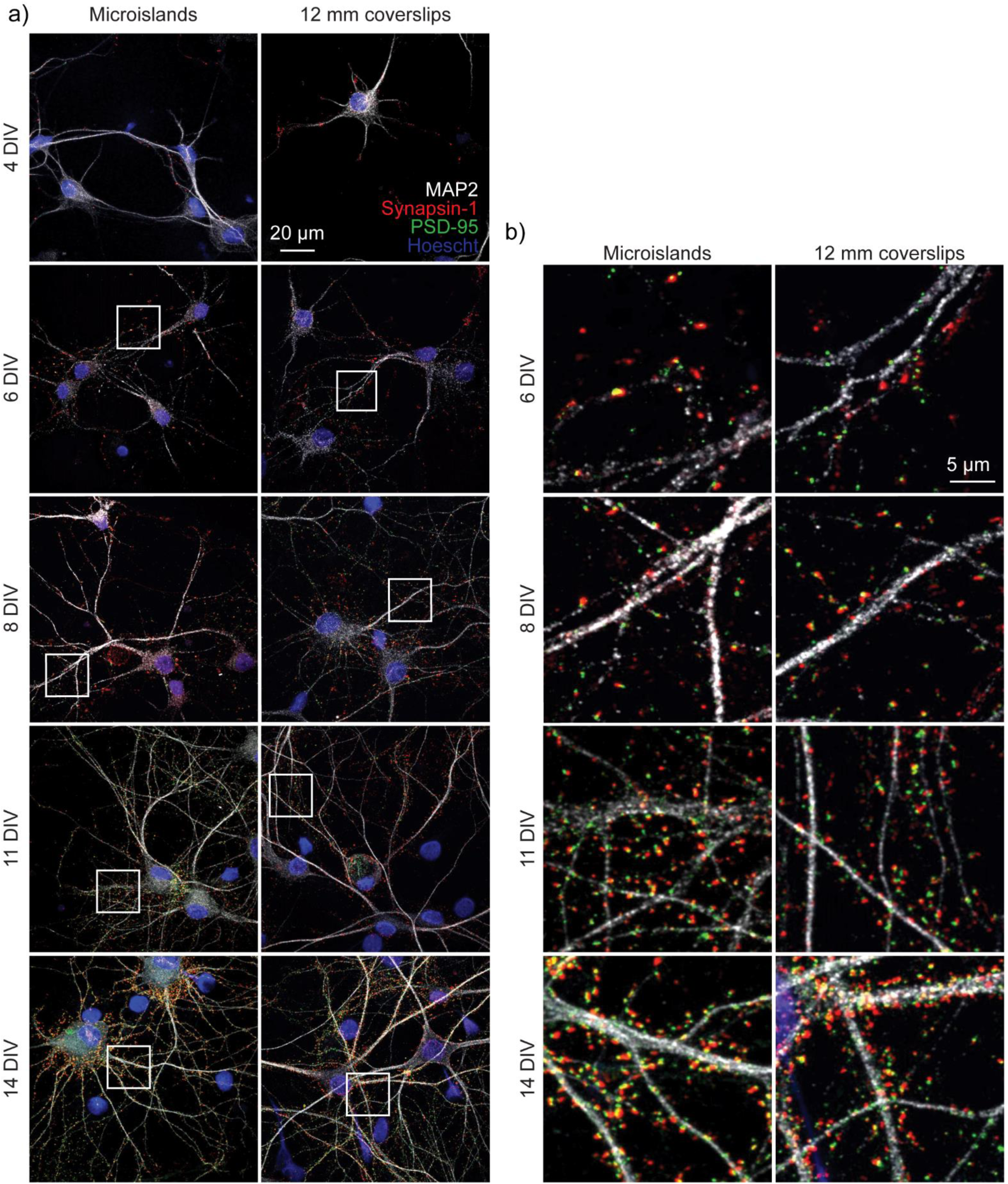
Immunohistochemistry of microislands and bulk (12 mm coverslip) culture. **a**) Confocal images of immunolabelled hippocampal cultures grown in microislands (left panels) or on 12 mm diameter coverglass (right panels) at different developmental stages (days *in vitro*, or DIV) as indicated. Dendrites were labelled with MAP2 (white), presynaptic terminals with Synapsin-1 (red), excitatory postsynapses with PSD-95 (green), and nuclei using Hoescht 33342 (blue). **b**) Zoomed in regions to show synaptic marker localization from images in (**a**) where areas are indicated by white boxes.

To assess the function of neurons within the entire microisland, we used the Optical Spike and Connectivity Assay, or OSCA, (Walker et al., 2020) which couples voltage imaging of the voltage sensitive fluorophore, BeRST 1, with a k-means clustering-based spike detection algorithms (Spike Connect). We previously showed that this technique quantifies action potential spiking patterns with comparable confidence to electrophysiological patch-clamp techniques (Walker et al., 2020). Using OSCA in microisland cultures, we find a striking similarity in firing frequency, inter-spike interval (ISI), and STTC recorded in microislands (**Figure 3**) to analogous data recorded previously in larger networks (Walker et al., 2020). As we discovered in larger, bulk dissociated hippocampal neuronal cultures, action potential frequency in microislands increases between 8 and 12 DIV, before decreasing back to the level initially observed at 8 DIV (**Figure 3a and b**). Moreover, like in bulk culture, inter-spike intervals for individual neurons in microisland culture at 8 DIV reveal a quasi-normal distribution spanning the range of possible inter-spike intervals (6 – 9000 ms), which, by 12 DIV, matured into preferred bands of spike timings centered at approximately 3 and 6 ln ISI (**Figure 3c and d**). Finally, neuronal connectivity increases across *in vitro* development. In microisland cultures, the spike-time tiling coefficient (STTC) values, a frequency-independent estimate of neuronal activity correlation and connectivity, show a stepwise increase throughout *in vitro* neuronal maturation (**Figure 3e,f**). These data closely mirror functional development data recorded with OSCA in bulk culture (Walker et al., 2020).

**Figure 3.**
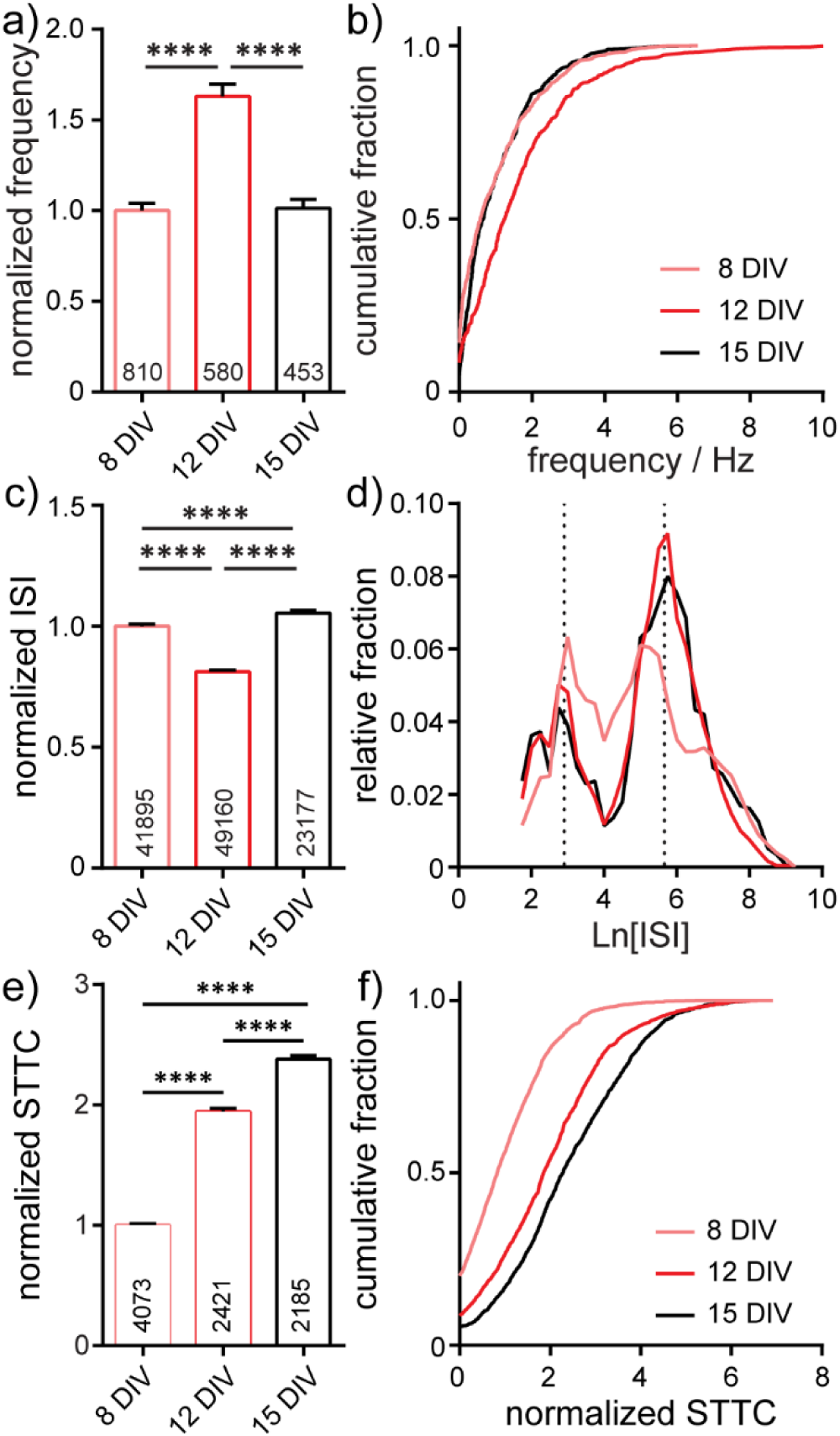
Characterization of neuronal activity in 650 × 120 μm microisland culture. Analyses of action potential frequency **(a,b)**, inter-spike interval, ISI, **(c,d)**, or spike timing tiling coefficient, STTC, **(e,f)** over developmental stages (8, 12, and 15 *days in vitro*; DIV) are summarized as a bar graph **(a,c,e)** and as a cumulative frequency plot **(b,f)**, or histogram **(d)**. Values on bar graphs indicate numbers of neurons **(a)**, pairs of spikes **(c)**, or pairs of neurons **(e)** analyzed for each condition. Data represent 4 biological replicates. Frequency, ISI, and STTC values are normalized to the 8 DIV value per biological replicate. Statistical tests are Kruskal-Wallis ANOVAs with multiple comparison tests to all groups. * = p <0.05, ** = p < 0.01, *** = p < 0.001, **** = p < 0.0001.

In summary, when evaluating synaptic marker expression (**Figure 2**) or voltage dynamics, including firing rate changes in development, ISI, and connectivity, as measured by STTC (**Figure 3**), we observe no differences in the *in vitro* development of microislands compared to bulk coverslip culture which has over 1000x larger surface area. Together these data suggest that voltage dynamics of hippocampal networks are unchanged by circuit size and support the use of patterned microislands as a model system to interrogate the wiring and connectivity of neuronal ensembles.

### Post hoc immunostaining enables cell-type specific analysis of firing frequency

The output of neural circuits is complex and depends on the inputs-to and responses-of large numbers of neurons made up of different cell types with a range of intrinsic properties. The neuronal networks of the hippocampus have been widely studied *in vivo*, owing to the roles of the hippocampus in learning and memory as well as its highly laminar structure. The principal excitatory neurons of the hippocampus, dentate granule cells (DGCs), CA3, and CA1, and the many subclasses of interneurons which innervate them, have their own molecular, morphological and functional characteristics (Wheeler et al., 2015). *In vitro*, without hippocampal architecture, assigning function to specific cell types is more challenging. Pairing two-cell electrophysiology with *post hoc* immunostaining has been applied to link cellular identity to functional output but is very low throughput (Williams et al., 2011). Here, to evaluate whether functional and molecular signatures of different cell types are maintained in microisland cultures, we paired OSCA with *post hoc* immunocytochemistry.

After imaging sessions with BeRST 1, we fixed the neurons and performed immunocytochemistry to assay for the presence of CaMKII, GABA, CTIP-2, and Prox1 (**Figure 4a**). Building on previous studies (Williams et al., 2011), we assigned cellular identities based on the combination of staining patterns. CaMKII-negative, GABA-positive were assigned as inhibitory cells (“inhib.” **Figure 4a and b**). CaMKII- and Prox1-positive cells were identified as dentate granule cells (“DGC”, **Figure 4a and c**) (Bagri et al., 2002); CaMKII- and CTIP-2-positive, Prox1-negative cells were identified as CA1 cells (“CA1”, **Figure 4a and c**) (Arlotta et al., 2005); cells which were positive for CaMKII, but negative for CTIP-2 or Prox1 were classified as CA3 cells (“CA3”, **Figure 4a and c**) (Williams et al., 2011; Evans et al., 2013); In this way, we could assign major classes of cells in the hippocampus: inhibitory, DGC, CA1, and CA3. Immunocytochemistry reveals that microislands (**Figure 4f and g**) have a similar cell-type composition to bulk cultures (**Figure 4d and e**). Both possess approximately equal proportions of principal excitatory neurons: DGCs account for 34% of identified cells in microislands and 22% for bulk; CA1 cells comprise 20% for microislands, 23% for bulk; and CA3 make up 30% for microislands and 39% for bulk. Inhibitory interneurons make up a smaller fraction of cells in both microislands (16% of 1,126 cells analyzed) and bulk culture (16% of 715 cells examined). There were instances where a microisland did not contain any inhibitory neurons. We analyzed microislands that did not contain any inhibitory neurons separately from those containing one or more interneurons and report on both groups below.

**Figure 4.**
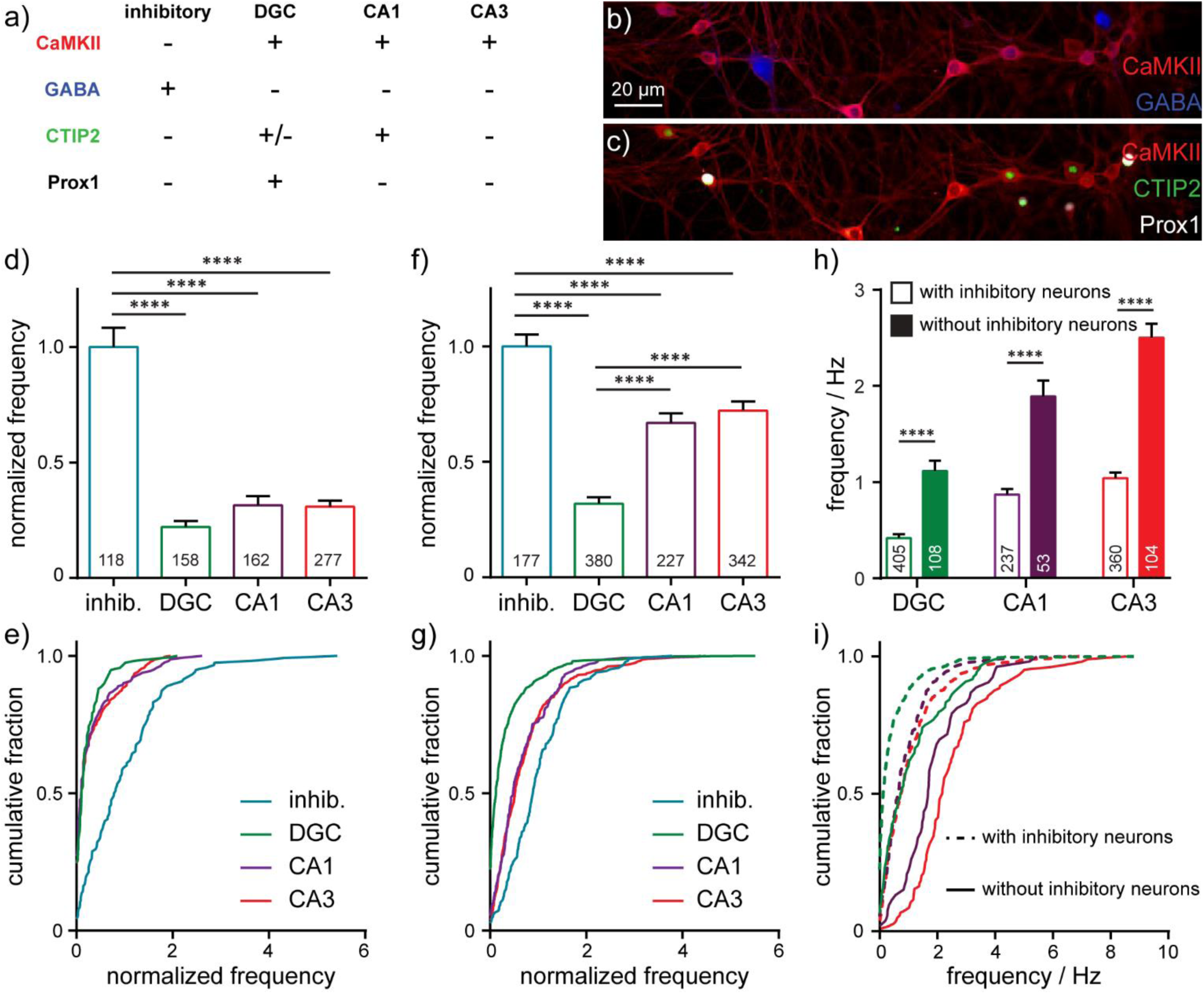
Cell-type dependent activity profiles interrogated by BeRST 1 voltage imaging. **(a)** Assignment of hippocampal cell type by post hoc analysis of cell type by immunofluorescence using anti-CaMKII, anti-GABA, anti-CTIP2, and anti-Prox1 antibodies. Images are of neurons fixed after functional imaging with BeRST 1 (as in **Figure 1**) and then stained for **(b)** CaMKII (red) and GABA (blue) or **(c)** CaMKII (red), CTIP2 (green), and Prox1 (white). Action potential frequencies for different cell types—inhibitory (“inhib.”, teal), dentate granule cells (“DGC”, green), CA1 (purple), and CA3 (red) pyramidal neurons—are summarized either as bar graphs for neurons grown **(d)** on 35 mm diameter coverslips and **(f)** in microislands containing at least 1 inhibitory neuron and as cumulative frequency plots for neurons **(e)** on coverslips or **(g)** in microislands. Numbers in bar graphs indicate the number of neurons analyzed across *n* = 5 (coverslips) or 11 (microislands) independent experiments (biological replicates). The difference in action potential frequency recorded from neurons in microislands with or without inhibitory neurons is summarized as **(h)** a bar graph or **(i)** cumulative frequency plot. Empty bars **(h)** and dashed lines **(i)** indicate values for microislands with at least 1 inhibitory neuron. Filled bar **(h)** and solid lines **(i)** indicate values for microislands without identifiable inhibitory neurons. Frequency, where indicated, is normalized to the frequency of inhibitory neurons measured for each biological replicate. Statistical tests are Kruskal-Wallis ANOVAs with multiple comparison tests to all groups **(d** and **f)** or Mann-Whitney t-tests for each cell type **(h)**. * = p <0.05, ** = p < 0.01, *** = p < 0.001, **** = p < 0.0001.

We find that interneurons have the highest mean firing frequency, whereas DGCs have the lowest mean firing frequency. CA1 and CA3 neuron sub-types exhibit intermediate firing rates which are not different from each other (**Figure. 4f-g**). Interestingly, approximately 25% of DGCs in microislands are quiescent during recordings, whereas almost all inhibitory and pyramidal neurons fire at least 1 action potential (**Figure 4g**). Action potential firing frequencies for different cell types were similar in microisland (**Figure 4f and g**) and bulk culture (**Figure 4d and e**).

In cases where no inhibitory neurons are present in a microisland, we observe mean firing frequency increases of ~2.5 fold across all excitatory neuron subtypes (**Figure 4h**) and the fraction of quiescent neurons drops to zero for all cell types including DGCs (**Figure 4i**) consistent with runaway recurrent excitation in the absence of inhibitory interneurons. Interestingly, even in the absence of inhibitory neurons, DGCs fired action potentials less frequently than pyramidal neurons, empirically demonstrating the intrinsic high threshold properties of DGCs (Spruston and Johnston, 1992; Penttonen et al., 1997; Nelson and Chris, 2006; Johnson-Venkatesh et al., 2015) even within a highly excited network (**Figure 4h and i**).

In summary, the activity we record for different hippocampal neuronal subtypes in both microislands and bulk culture are consistent with reported values for excitatory and inhibitory hippocampal neurons in intact preparations: inhibitory neurons spike more often, and excitatory neurons show lower levels of activity, with DGCs firing significantly less frequently that pyramidal neurons (Spruston and Johnston, 1992; Nelson and Chris, 2006). These data indicate that several gross functional features of hippocampal cell types are conserved in dissociated microisland systems.

### Principal excitatory neurons preferentially wire in microislands

*In vivo*, the hippocampus is a laminar brain structure with stereotyped, mostly unidirectional circuitry, with information flowing from DGCs to CA3 and then to CA1 neurons, before leaving the hippocampus. During dissociation of the hippocampus in primary culture, the laminar structure and associated spatial connectivity of the hippocampus is initially lost. However, even reductionist systems like dissociated hippocampal neuron culture recapitulate aspects of the astonishing specificity of neuronal wiring found in intact brains (Williams et al., 2011). Given that the identity and functional signatures of neuronal subtypes are maintained in microcircuits, we decided to examine whether any elements of native functional connectivity were preferentially formed and could be detected using non-invasive and higher throughput voltage imaging techniques.

To determine functional connectivity we looked at the timing of action potentials between every possible connection, hypothesizing that if neuron A is monosynaptically connected to neuron B, we would expect a delay in action potential firing in neuron B relative to action potential firing in neuron A. To capture the length of this delay we examined cross-correlation timings between spikes in pairs of neurons across all of our imaging bouts, revealing patterns in the timing of firing between pairs of neurons (**Figure 5a**). Most often, spikes between neurons occurred co-incidently, indicating the highly interconnected nature of neurons in culture. A second peak in spike timing delays (lag time) was observed between 8 to 14 ms, corresponding to monosynaptically driven functional connectivity. A third peak, observed between 18 to 26 ms, likely represents disynaptic connections.

**Figure 5.**
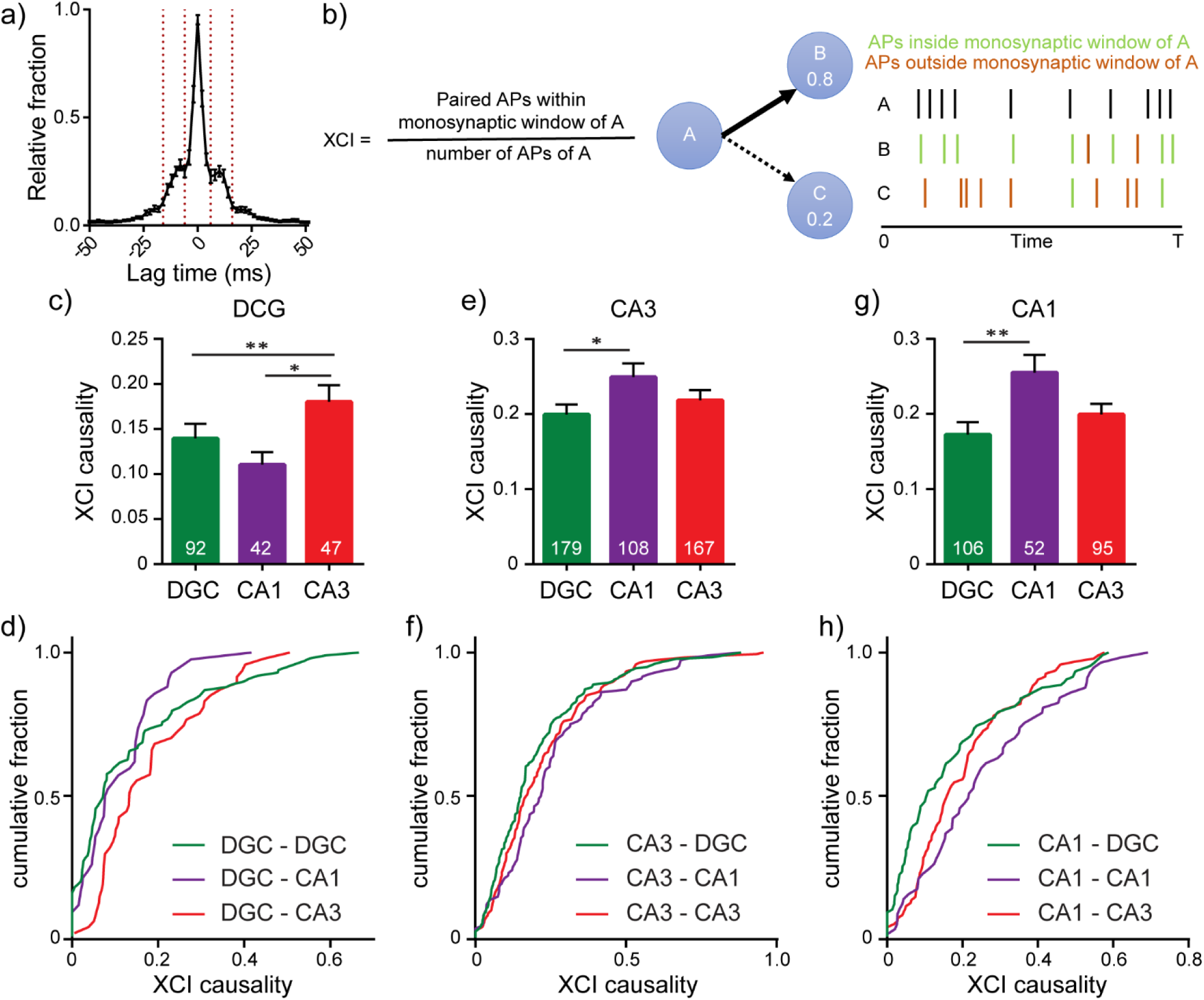
Interrogating synaptic specificity in neuronal networks using voltage imaging. **a)** Histogram of cross-correlation lag times between spikes in microisland cultures. Vertical lines highlight the 8 – 14 ms window we identified as a monosynaptic lag. **b**) Schematic detailing XCI causality, a score of the strength of connection between neurons determined by comparing firing patterns of a template neuron with all excitatory neurons within the circuit. Bar plots of XCI causality and associated cumulative frequency plots showing connectivity strength between template neurons (indicated at the top of the graph) and other excitatory neurons in the same network at 15 DIV, where the template neurons are either DGCs (**c-d**) CA3 neurons (**d-f**) or CA1 neurons (**g-h**). n indicated on graphs are numbers of neuron pairs analyzed. Data recorded from 42 microislands. Biological n is 4 experiments. Statistical tests are Kruskal-Wallis ANOVAs with multiple comparisons tests to all groups. * = p <0.05, ** = p < 0.01.

Using the monosynaptic window of 8 to 14 ms, we generated a cross-correlation index (XCI) metric to quantify the likelihood of a neuron causing an action potential in any other neuron via a monosynaptic connection, called XCI causality. For given pair of cells, for example, cell A and B, we define XCI as the number of spikes in B occurring within the monosynaptic time window (8 to 14 ms) of A divided by the total number of spikes produced by cell A (**Figure 5b**). XCI is therefore a metric of the likelihood that a spike in cell A triggers a spike in cell B. It has the typical bounds: 1.0 indicates that all of the spikes in cell A result in spikes in cell B within a monosynaptic time window, and 0 indicates that none of the spikes in cell A trigger a spike in cell B. By calculating the XCI for each pair of neurons in the microisland and complementing this with cellular identity information, determined *post hoc* by immunostaining, we developed probability maps for neuronal networks. We examined the likelihood of each excitatory neuron sub-type (DG, CA1, CA3) to trigger a spike in a monosynaptic connection (**Figure 5c-h**). XCI values cover a broad range across all connection types. However, each triggering neuronal subtype, possesses a preferential partner. We find that DGCs (**Figure. 5c and d**) are more likely to trigger firing in CA3 neurons (XCI = 0.22) rather than in either CA1 neurons (XCI = 0.11) or in other DGCs (XCI = 0.14), suggesting that DGCs form the strongest functional connections with CA3 neurons. This agrees with hippocampal wiring *in vivo* and previous studies that establish DGC to CA3 synapse specificity in culture (Williams et al., 2011). However, preferential wiring between other cell types *in vitro* has not been reported. Here, profiling connections made from CA3 neurons, we find that CA3 neurons are more likely to trigger action potentials in CA1 neurons compared to DGCs (XCI = 0.25 vs 0.2, p < 0.05, **Figure 5e and f**), in agreement with native wiring *in vivo*. CA1 neurons in hippocampal culture lack their endogenous postsynaptic partners in the entorhinal cortex and subiculum and instead preferentially wire recurrently to trigger spikes in other CA1 neurons (XCI = 0.26, **Figure 5g and h**). In summary, quantification of functional connectivity between the three principal excitatory neuronal subtypes reveal that, within a backdrop of high interconnectivity, preferential wiring is maintained *in vitro*. Using OSCA and patterned microislands enables the simultaneous interrogation of all classes of connections between cell types, limited only by the number of markers used to define cellular identity.

## Discussion

In this paper we present a method to interrogate neuronal microcircuitry *in vitro*. To achieve this, we couple voltage imaging-based action potential detection with neuronal subtype identification using immunocytochemistry in the context of patterned microislands. The flow of neural information throughout these circuits was elucidated by simply following sequences of action potentials in real time and was made possible by imaging every single neuron belonging to the network.

Reducing the network size was a key advance for this work as it removed inputs to and outputs from the recording area that we simply could not trace to their source or termination in larger, bulk cultures. Removing this outside influence reduced noise levels and meant that all action potentials recorded had their origin and propagation within our window of observation. However, in undertaking this reductionist view it was essential to characterize how low neuronal numbers present in microislands might alter circuit activity. In fact, we found that shrinking network size to only 10-25 cells had no effect on the functional behavior of the circuit as long as at least one inhibitory neuron was present: the evolution of firing frequency, inter-spike intervals, and connectivity (STTC) across maturation remain indistinguishable between microislands and bulk cultures over 1000 times larger. Furthermore, activity rates measured in neuronal subtypes (DGC, CA3, CA1, Inhibitory) were also conserved across the two culture sizes. This is perhaps not surprising considering the plastic nature of neurons and their ability to undergo homeostatic alterations to reach a set point of functional activity. However, neurons and circuits can only remodel within limits. We found that if a circuit contained zero inhibitory neurons, activity patterns and firing frequencies across all cell types increase due to recurrent excitation (**Figure 4h and i**).

Our ability to record action potential firing in real time opened the possibility of converting the precise timings of monosynaptic communication to infer functional connectivity. We assessed whether preferential wiring between hippocampal excitatory subtypes is conserved *in vitro*. Previous work showed that DGCs preferentially synapse onto CA3 pyramidal neurons in dissociated culture (Williams et al., 2011). This was achieved using an impressive and painstaking paired electrophysiological technique where neurotransmitter release was triggered via an electrode in DGCs and excitatory postsynaptic potentials were recorded in putative partners, one neuron at a time. While yielding important information on DGC partners, this approach, similar to connectomics, outlines whether a connection exists, not whether it is actually activated during nascent circuit activity. The approach used here, measuring the likelihood of action potential firing in one neuron to trigger action potential firing in a downstream neuron in a spontaneously active network, however, is a direct measure of information propagation and holistically sums the effects of synaptic strength, dendritic integration, and intrinsic excitability on neuronal output. Additionally, because the entire neuronal network is sampled simultaneously, this strategy has the power to go further, revealing all functional connections between all cell types. Indeed, we show that in dissociated microisland culture, DGCs preferentially trigger action potential firing in CA3 neurons, while CA3 neurons preferentially trigger action potential firing in CA1 neurons, matching hippocampal circuitry *in vivo*. CA1 neurons, lacking their endogenous postsynaptic partners, preferentially trigger firing in other CA1 neurons. Because we quantify the functional connectivity between every neuron pair, we also capture the noise in the system, and find that, in addition to preferential connectivity, there is a high level of general connectivity between all excitatory subtypes. In summary, these data show that hippocampal circuitry is broadly recapitulated *in vitro*, indicating that hippocampal microisland models are a valid model for studying hippocampal circuitry *in vitro*.

### Limitations and future considerations

This study occupies an important gap in the field of neural circuits: the ability to provide a functional readout of connectivity on time and spatial scales that can resolve neuron-neuron interactions for identified cells, and to achieve this for many connections simultaneously. This combination of detail and throughput cannot currently be achieved with any single calcium imaging, multielectrode array (MEA), or electrophysiological modality. However, these experiments are bound by several important considerations. First, the studies described here were carried out on small numbers of connected neurons (<25).

Ideally, experiments of this sort would be carried out on even larger ensembles of neurons. The low neuron numbers are not a limitation of the microisland patterning, but of the camera we employed in this study. To achieve a framerate of 500 Hz and enable the reconstruction of precise spike timing across different cell types, we had to scale down the size of the field of view captured by the sCMOS camera. As new generations of sCMOS and EMCCD cameras emerge, this will improve our ability to record from larger numbers of neurons. Additionally, the use of high NA, low magnification objective lenses and custom microscopes will enable recording from even larger regions (Werley et al., 2017).

Second, in this study we relied on four markers (anti-GABA, -CaMKII, -CTIP2, and -Prox1) to classify four different cell types in hippocampal culture. The methods descried here could be combined with additional, or complementary, *post hoc* analysis to classify cell types. For example, analysis of RNA transcript levels would provide a simultaneously more expansive and fine-grained view of cellular identity, which could be paired with existing and emerging cellular inventories of brain regions and cell types (Ecker et al., 2017).

Third, the spike timings in this study were determined from recurrent, spontaneous activity. Stimulation or inhibition of specific cells or cell types with opsins (Deisseroth, 2015) or pharmacology, or activation/blockade of specific channels via photoswitchable ligands (Hüll et al., 2018; Tochitsky et al., 2018), would provide opportunities to explore dynamic remodeling of neuronal networks. Because the presence of just a single inhibitory neuron in patterned microisland culture profoundly alters spiking frequency (**Figure 4**), the dynamic inclusion or removal of targeted neuron classes could be a powerful method for studying neuronal and circuit plasticity.

Fourth, the neuronal ensembles under observation here lack the context of an intact preparation. Despite this fact, data we present here points towards hippocampal microisland cultures recapitulating the specific wiring of native circuitry found *in vivo*, particularly along the DG to CA3 and CA3 to CA1 axis. This is supported by previous studies highlighting specific DG to CA3 connections in hippocampal cultures (Williams et al., 2011). This makes patterned microislands, coupled to voltage imaging with OSCA, a reasonable starting point to look for the molecules the underlie this remarkable specificity, for example Cad9, as previously identified (Williams et al., 2011), or to explore other molecules that may alter or modulate these activity patterns.

Finally, the neurons used here were from healthy animals. The methods we describe are amenable to the use of neurons isolated from disease models, or differentiated from patient-derived stem cells. We envision a role for the methods described here in unraveling mechanisms underlying circuit breakdown in disease. Circuit dysfunction is key phenotype in many neurological and neurodegenerative diseases. Often, circuit dysfunction accompanies the loss of certain neuronal subtypes, for example dopaminergic substantia nigra neurons in Parkinson’s Disease and hippocampal CA1 neurons in Alzheimer’s disease, in a process termed selective vulnerability. Using this technique, it should be possible to track how the loss of specific cell types affects the ongoing activity and function of their circuits. Here the adaptability of this assay is a key advantage as many different circuits can be assessed simply by changing the brain regions dissected.

## Acknowledgements

We acknowledge support from the NIH (R01NS098088). BKR was supported in part by an NIH Training Grant (T32GM066698). Confocal microscopy was performed at the CRL Molecular Imaging Center at UC Berkeley, supported by the Helen Wills Neuroscience Institute.

